# Genomic analyses of recently emerging clades of mpox virus reveal gene deletion and single nucleotide polymorphisms that correlate with altered virulence and transmission

**DOI:** 10.1101/2024.09.24.614696

**Authors:** Ankeet Kumar, Pratima Jhanwar, B Roohani, Amulya Gulati, Utpal Tatu

**Affiliations:** Department of Biochemistry, Division of Biological Sciences, Indian Institute of Science, Bangalore, India

**Keywords:** Mpox virus, Mutational analysis, Clade Ib, Complement control protein, Single nucleotide polymorphisms

## Abstract

Mpox virus (MPXV) has consistently caused human infections since the first reported case in 1970, with the initial outbreaks primarily attributed to sporadic zoonotic transmissions. In recent years, an increase in human-to-human transmission has been observed, particularly during the 2022 outbreak caused by Clade IIb, which was declared a public health emergency by WHO. In 2024, the emergence of Clade Ib from Africa raised global concern once again. While several studies have provided valuable insights into the differences among MPXV clades, research on Clade Ib remains limited. In this study, we have conducted a comprehensive comparative genome sequence analysis and identified unique features across MPXV clade sequences. We report a ∼1141 bp long novel deletion resulting in loss of the complement control protein (CCP) in Clade Ib sequences, a deletion earlier exclusively reported in Clade II sequences. Additionally, B22R, a crucial host receptor-binding protein involved in viral entry and known for its high immunogenicity, shows clade-specific amino acid changes across three MPXV clades. Moreover, multiple extragenic mutations were identified in the 5’ UTR of several genes, which may impact gene transcription. Other frequently mutated proteins are linked to immune evasion, translation, and viral entry and exit. The above results and APOBEC signatures associated with sustained human-to-human transmission highlight the virus’s potential for rapid adaptation, underscoring the need for vigilance against reverse zoonosis and the risk of spillover to new hosts.

## Introduction

Mpox, formerly known as monkeypox, is a zoonotic infection caused by the mpox virus (MPXV), a double-stranded DNA virus of the genus *Orthopoxvirus* and family Poxviridae. Phylogenetically, MPXV is closely related to other orthopoxviruses such as vaccinia, cowpox, and variola viruses, sharing significant genetic similarities [1]. The disease is characterised by skin lesions, and other common symptoms of mpox in humans include fever, chills, muscle ache, fatigue, and enlarged lymph nodes [2]. Traditionally thought to be transmitted through direct contact with infected individuals, including their rash, scabs, and bodily fluids [2, 3], mpox is now also suspected to spread via indirect contact, such as aerosols and contaminated surfaces [4].

MPXV was first identified and isolated in 1958 in a primate research facility in Denmark from monkeys that had been transported from Singapore for research purposes [5]. Although it was initially discovered in monkeys, the primary reservoir of the virus is believed to be small mammals like rodents [6]. The first reported human case of mpox was a nine-month-old boy in the Democratic Republic of Congo (DRC) in 1970 [3, 7]. Mpox was endemic to Africa for a long time, with cases primarily confined to the DRC, Nigeria and neighbouring countries [8]. In 2003, the first cases outside Africa were reported in pet prairie dogs in the United States; these pets shared bedding and caging with a shipment of infected small mammals from West Africa [9]. This outbreak resulted in 47 confirmed human cases, with no evidence of human-to-human transmission [10]. Since then, various small outbreaks have occurred, predominantly in African countries [11]. Following a major international outbreak in 2022 caused by Clade IIb, a different clade of MPXV (Clade Ib) has been spreading outside of Africa in 2024, with recent reports of human cases in Sweden (August 15, 2024), and Thailand (August 22, 2024). This resurgence highlights the public health challenges associated with the virus and the need for continued vigilance and international cooperation in controlling this menace.

The genome of orthopoxviruses is linear, double-stranded DNA (dsDNA) and ranges from 170 to 250 kb in size [12]. The genome is flanked by inverted terminal repeat (ITR) sequences that form covalently closed hairpin structures at the termini [13]. The number of genes in orthopoxviruses ranges from around 160 genes in the taterapox virus to about 210 genes in certain strains of the cowpox virus [14]. The MPXV genome is around 196.8 kb in size and encodes approximately 180 proteins [15]. Two distinct clades of MPXV, the Congo Basin or Central African Clade and the West African Clade (later renamed as Clade I and Clade II, respectively), emerged early in the history of the MPXV [16]. Over time, genome diversification has led to the formation of more clades, resulting in Clade I being divided into Clades Ia and Ib, and Clade II into Clades IIa and IIb.

To investigate the emergence of sustained human-to-human transmission in novel MPXV variants (Clades Ib and IIb), and gain insights into the molecular differences among clades, we conducted a comparative genomic analysis across MPXV clades, utilising complete genome sequences from public databases. Our analysis revealed several mutation types, including gene truncations, frameshifts, nucleotide insertions and deletions, and loss of stop codons, across 164 viral genes. A total of 41 non-synonymous amino acid changes were identified across all four clades, distributed among 26 proteins. Notably, around 19% of these changes occurred in the B22R family protein encoded by MPXV_gp174, followed by the DNA-dependent RNA polymerase subunit rpo147 (MPXV_gp0085) and ribonucleoside-diphosphate reductase 2 (MPXV_gp060). Additionally, we identified 87 amino acid changes unique to three clades (Clade Ib, Clade IIa, Clade IIb), with the highest number of mutations in Clade IIb (43), followed by Clade IIa (26), and Clade Ib (18), distributed across 33, 22, and 16 proteins, respectively. These high-frequency mutations primarily affect distinct proteins, indicating possible biological differences among the clades. Some of these mutations, based on MutPred2 and PolyPhen2 predictions, may influence protein function, although further experimental validation is needed. Nonetheless, they hold potential as markers for rapid, clade-specific tests, enhancing surveillance and aiding in more efficient resource allocation. We also observed the deletion of the complement control protein (CCP) in Clade Ib, previously reported only in Clade II (Clades IIa and IIb). This deletion indicates a possible functional adaptation that may influence the virus’s interaction with the host immune system. Furthermore, APOBEC signatures revealed markers of sustained human-to-human transmission in recent variants. The above findings highlight critical differences in the clade genomes with distinct mutational repertoires. These insights are essential for understanding the virus pathogenesis and transmission dynamics, which will help enhance preparedness for future outbreaks and inform preventive measures, particularly in the context of increasing transboundary viral spread and sustained human-to-human transmission.

## Methods

### Data Procurement and Curation

Complete and near-complete MPXV whole genome sequences, along with their accompanying metadata were obtained from two databases - Nextstrain (Dated: 22-08-2024) [17] and Global Initiative on Sharing All Influenza Data, GISAID (Dated: 28-08-2024) [18]. The data was curated manually to remove the sequences that contained >600 N characters, followed by removing sequences that lacked complete metadata. The Clade Ib sequences from the recent outbreak (2024) were obtained from GISAID, which included cases outside Africa, specifically from Thailand (EPIISL19350788).

### Alignment and Phylogeny

The MPXV genomes were aligned using MAFFT v7.490 (command line) [19]. The alignment files were renamed to include information such as accession number, host, country, collection year, and clade. The aligned sequences were then used to generate a maximum likelihood phylogenetic tree using IQ-TREE v2.3.5 for Linux [20], employing the model suggested by minimum BIC and AIC scores. The resulting tree file was visualised and annotated using iTOL (webserver) [21].

### Single Nucleotide Polymorphism Analysis

Nucleotide variations were identified across all complete and near-complete MPXV genomes relative to the NCBI reference sequence (NC_003310.1), using Nucmer v3.1 [22]. Nucmer is a command-line tool which performs pairwise sequence alignment between reference and query sequences, producing a table with details on nucleotide variations and their coordinates. This nucleotide variation data was analysed using an R script, which was originally developed for severe acute respiratory syndrome coronavirus 2, later improved for other viruses [23–25]. This script was further modified to meet the specific requirements of the MPXV genome. The previous version of the script, which was limited to handling sequences with positive polarity only, was modified to accommodate the ambipolar nature of the MPXV genome. This modified script is provided as Supplementary File 1. Another R script that was developed to identify non-overlapping, unique, and high-frequency (>90%) mutations for the MPXV clades is available in Supplementary File 2. A general feature format 3 (gff3) file with the coordinates of coding regions in the MPXV genome is presented in Supplementary file 3.

### Analysing the Impact of Mutations

MutPred2 and PolyPhen2 were employed to assess the impact of frequently occurring mutations on protein structure and function [26–28]. PolyPhen2 evaluates the overall effect of mutations on protein function, while MutPred2 provides additional insights into how these mutations might alter specific molecular mechanisms.

### Tools used

Other tools used are Simplot++ [29], Highlighter [30], DnaSP v6.1 [31], NCBI MSA Viewer 1.25.0, Excel, Python (Pandas, Matplotlib), R v4.1 (ggplot2, dplyr) and Inkscape.

## Results

### Phylogenetic and DNA Polymorphism Analysis Reveals Clade IIb as the Most Genetically Diverse MPXV Clade

Orthopoxviruses have periodically caused outbreaks, and the recent global emergence of these outbreaks has prompted the study of the evolutionary relationships among these viruses. Lately, MPXV has particularly raised concerns due to its increased transmission outside of Africa and its potential for human-to-human transmission. To examine the relationship of MPXV with other orthopoxviruses, a whole genome-based neighbour-joining (NJ) phylogeny of twelve orthopoxviruses was generated, using Yaba monkey tumour virus as an outgroup. As shown in Figure 1A, the phylogenetic tree of orthopoxviruses divides into two main branches. The first branch contains three viruses: raccoonpox virus, volepox virus, and skunkpox virus, which are prevalent in North America. The other branch harbours nine viruses: Akhmeta, Abatino, ectromelia, cowpox, taterapox, camelpox, vaccinia and MPXV (NC003310), which are found in Africa and Eurasia. MPXV is most-closely related to vaccinia virus, as they share the same phylogenetic node, indicating a common evolutionary ancestor.

**Figure 1.**
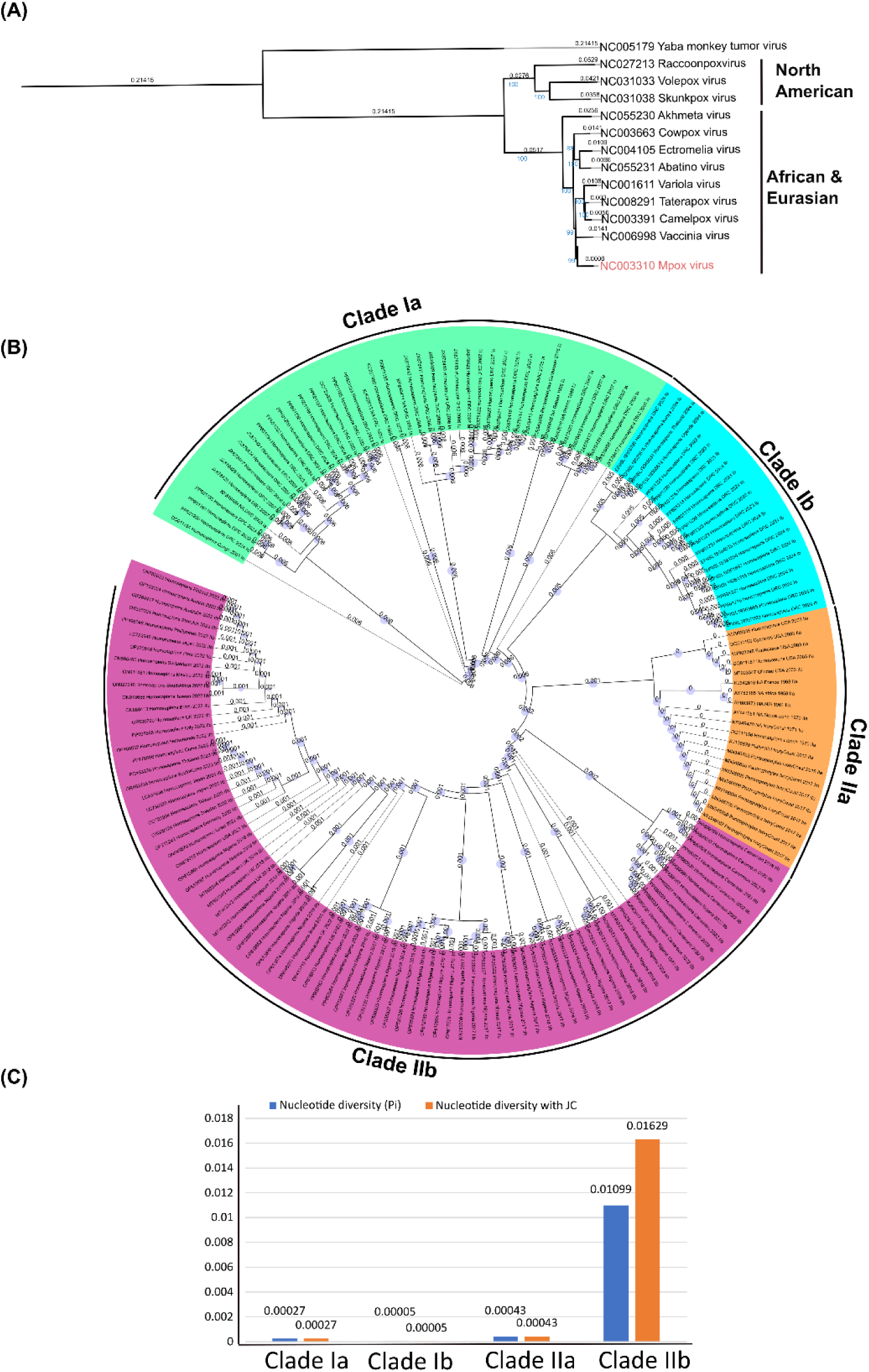
Phylogenetic analysis of MPXV clades and other orthopoxviruses. (A) A neighbour-joining (NJ) phylogenetic tree representing the relationship among twelve orthopoxviruses with Yaba monkey tumour virus as an outgroup. The distance values are shown above each branch in black, and bootstrap values in blue. (B) Phylogenetic distribution of MPXV whole genome sequences. A circular representation of the phylogenetic tree of all the sequences retrieved from Nextstrain and GISAID. Clade Ia, Ib, IIa, and IIb sequences are shown in green, blue, orange, and pink, respectively. (C) Nucleotide diversity in MPXV clades. Nucleotide diversity (π) values and nucleotide diversity corrected by the Jukes and Cantor (JC) model were calculated using the DNA polymorphism module of the DnaSP tool.

Figure S1A illustrates MPXV outbreaks since its discovery, with increased human-to-human transmission observed in recent years. Region-wise data on human cases and deaths from mpox are presented in Figure S1B. To examine the evolutionary relationships among these clades, a maximum-likelihood phylogeny based on 166 MPXV sequences isolated from six hosts (*Homo sapiens*, *Pan troglodytes*, *Gliridae*, *Funisciurus*, *Cynomys*, and *Platyrrhini*) is presented in Figure 1B. The phylogeny reveals the presence of four distinct clades - Clade Ia, Ib, IIa, and IIb, comprising sequences from 35 countries. Clade Ia sequences in our dataset are mostly from isolates from the DRC. Whereas, Clade Ib sequences, which are linked to the current (2024) outbreak, have been isolated from multiple regions, including countries surrounding the DRC, as well as in Asia and Europe. The sequences EPIISL19345034 from Thailand cluster on the same branch as EPIISL19350788 from the DRC. Clade IIa has been endemic to Western Africa, and its offshoot, Clade IIb, was responsible for the major global outbreak in 2022, affecting various countries and resulting in various lineages.

Figure 1C illustrates the nucleotide diversity for MPXV clades; Clade IIb shows the highest nucleotide diversity, followed by Clades IIa, Ia and Ib. The genome identity and similarity among MPXV clades and other orthopoxviruses are provided in Table S1. These observations highlight the evolutionary potential of MPXV clades, underscoring the need for enhanced surveillance and genome sequencing to uncover reservoirs, evolutionary patterns, and transmission dynamics.

### Extragenic Mutations Predominate Across All MPXV Clades

To understand the rapid emergence of new variants of MPXV, we analysed the mutational patterns among available MPXV sequences. Figure 2A shows the normalised average mutation count for each clade. Clade IIb, responsible for the 2022 outbreak, exhibits the highest average mutation count of 1387, whereas Clade IIa has 1292 average mutations. Clade Ia shows the lowest mutation count with an average of 300, and Clade Ib shows 329 mutations on average. Figure 2B shows the number of mutated proteins out of 180 annotated proteins. The highest number of mutated proteins are in Clade IIa, followed by Clades IIb, Ib, and Ia. This result highlights that Clades Ib and IIb, which are associated with more recent outbreaks, exhibit a greater degree of mutations, suggesting that these clades may be undergoing more rapid evolutionary changes.

**Figure 2.**
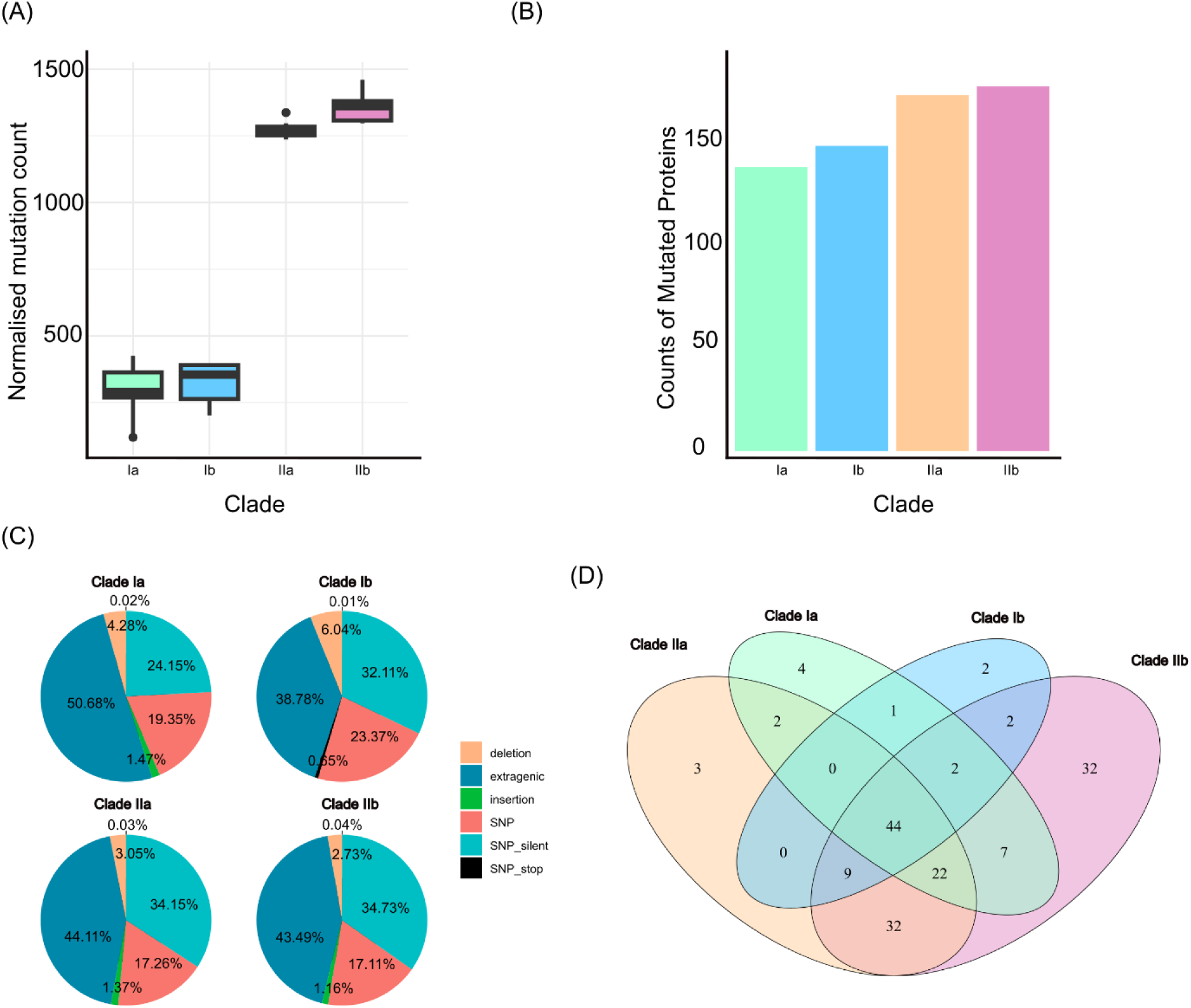
Clade-wise distribution of mutations among MPXV clades. (A) Mutation count normalised by the number of samples for each MPXV clade. (B) A graphical representation of the number of mutated proteins in all the clades depicted by the different coloured bars. Here, Clade Ia is shown in green, Ib in blue, IIa in orange, and IIb in pink. (C) A pie chart illustrates the different types of mutations i.e., extragenic, SNP, SNP-silent, deletion, insertion, SNP-stop mutations for each clade. (D) A Venn diagram illustrating the distribution of mutated proteins across different clades. Each clade is represented by an ellipse, with the number of shared and uniquely mutated proteins shown in the overlapping and non-overlapping regions of the ellipses, respectively.

Figure 2C presents a pie chart illustrating the variation in the percentages of mutation classes across the clades. Extragenic mutations are the most prevalent, ranging from approximately 39% to 51%, followed by synonymous and non-synonymous mutations, with deletions and insertions being the least prevalent among the mutation classes, and SNP-stop mutations accounting for less than 1% of the mutation classes. Figure 2D shows the distribution of mutated proteins across the MPXV clades. A total of 44 proteins are commonly mutated in all the clades, while the others are shared between one or more clades. The highest number of uniquely mutated proteins are found in Clade IIb (32), followed by Clades Ia (4), IIa (3) and Ib (2). The predominance of extragenic mutations across all the clades suggests that the regulatory regions play a crucial role in MPXV evolution.

### APOBEC Signatures Highlight Human Adaptation in Clades Ib and IIb

MPXV is known to infect various mammalian hosts, and as it adapts to these hosts, the virus undergoes multiple nucleotide-level changes. For example, dominant APOBEC (Apolipoprotein B mRNA Editing Catalytic Polypeptide-like) enzyme markers have been reported in Clade IIb [32]. We investigated the presence of APOBEC signatures in Clade Ib and compared them to those found in other MPXV clades. Figure 3A shows the trend of the top ten nucleotide changes across the MPXV clades, with cytosine to thymine (C>T) being the most dominant substitution in all clades. This is followed by guanine to adenine (G>A), while A>G is also notable. The trend of other transitions varies across the clades. The comparison of transitions and transversions across the complete genomes of four MPXV clades is shown in Figure S2. The increasing activity of APOBEC in novel variants suggests sustained human-to-human transmission.

**Figure 3.**
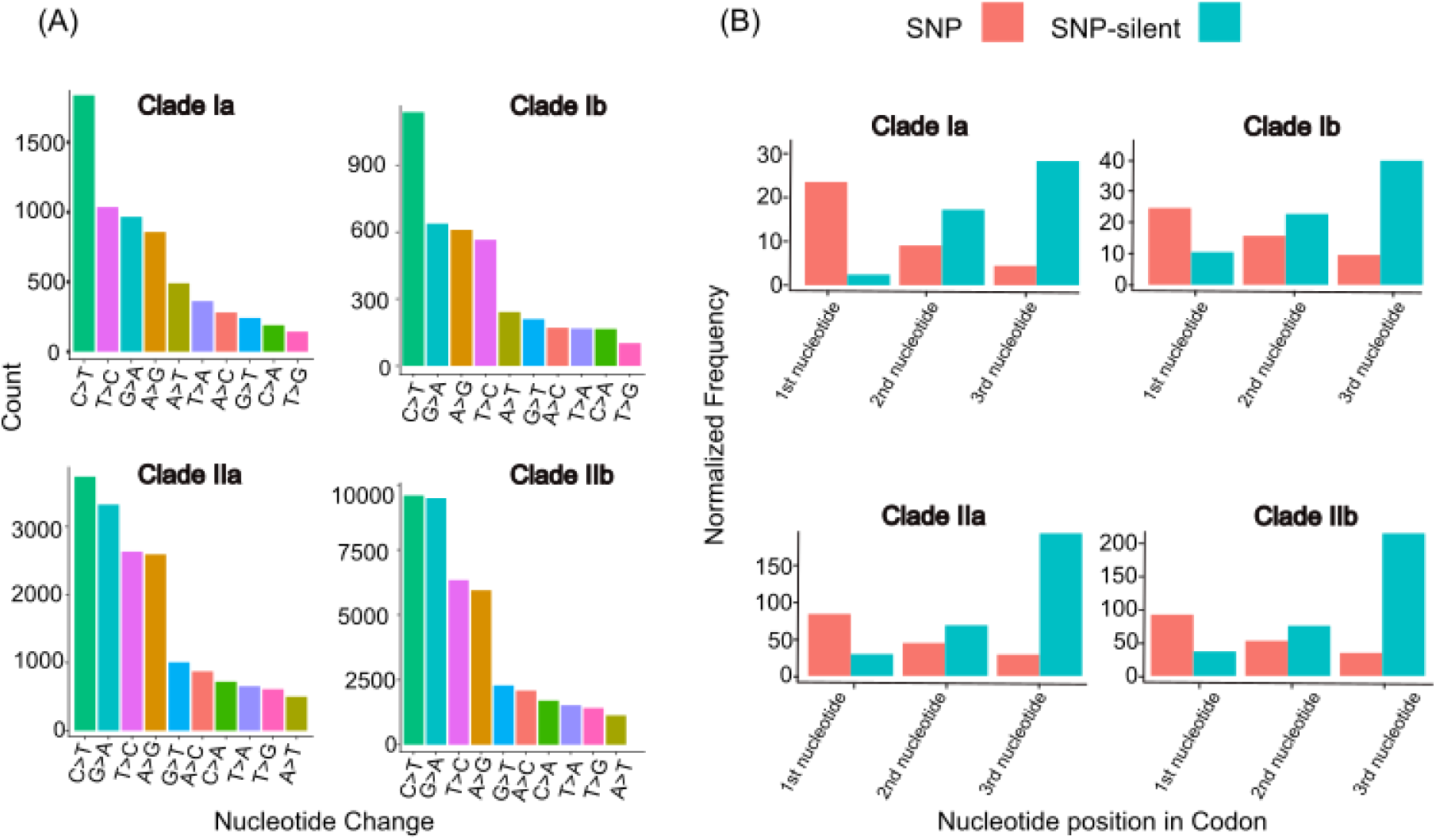
Nucleotide transitions and nucleotide position in codon in MPXV clades. (A) Representation of transitions and transversions among Clades Ia, Ib, IIa, and IIb in the form of histograms. X-axis shows different nucleotide changes depicted by different coloured bars, and their counts on the Y-axis. (B) Comparison of frequency (normalised to sample count) of SNP versus SNP-silent mutations resulting from nucleotide positions within a codon across the clades.

To investigate the selection pressures acting on specific nucleotide positions within codons, we conducted a comprehensive analysis across MPXV clades. Our findings reveal that more than 98% of both non-synonymous and synonymous mutations in MPXV arise from single-nucleotide changes (data not shown). Figure 3B demonstrates the distribution of nucleotide positions within codons that contribute to SNP-silent (synonymous) and SNP (non-synonymous) mutations across the clades. In Clade Ia, most non-synonymous changes are driven by alteration in the first nucleotide of the codon. In contrast, changes in both first and second nucleotide positions play significant roles in driving amino acid substitutions in other clades. Additionally, we observe high synonymous mutations in Clade II sequences which are mostly driven by change in the third nucleotide of the codon. These findings highlight the varying evolutionary pressures and codon usage patterns that may shape the adaptive strategies of MPXV.

### Genome Comparison Reveals Gaps in MPXV Clades and a Novel Deletion of CCP Protein in Clade Ib

Our previous results (Figure 2C) reveal that the mutation repertoire across all clades is predominantly composed of extragenic mutations, followed by SNP-silent, SNP, and frameshift mutations. To examine whether the emerging clades exhibit gaps or deletions in their genome, we compared the complete genome sequences of MPXV clades, which revealed gaps in various regions of the genome of different clades. Figure 4 depicts plots of pairwise comparisons among different MPXV clades relative to the MPXV reference genome (NC_003310), highlighting variations in the genome’s terminal regions while the central regions remain primarily conserved. A comparison of Clade Ia (NC_003310) versus Clade Ib reveals a gap of 1141 base pairs (bp), with ORF analysis indicating the deletion of a complement control protein (CCP) in the Clade Ib genome. In Clade IIa, there are three major gaps; the gap affecting CCP is 1953 bp long, and the other two gaps are present in the extragenic regions. In Clade IIb, four gaps are apparent, where a deletion of a 1953 bp region affects CCP, while the others are present in the extragenic regions. Other small gaps and insertions were also found in genes across these clades (data not shown). Figure S3 presents a multiple sequence alignment of Clade Ib sequences, demonstrating the conservation of CCP deletion across the sequences. Previous studies have reported the deletion of this gene only in Clade II (Clades IIa and IIb) sequences.

**Figure 4.**
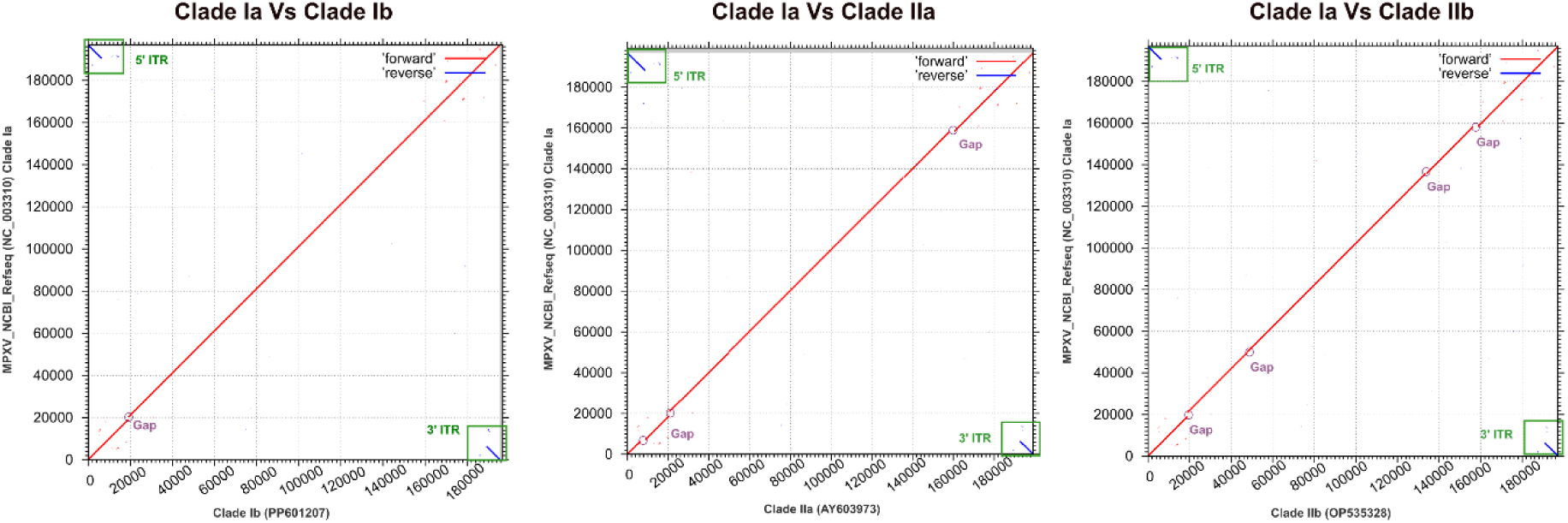
LAST hits plots showing sequence conservation and gaps across three clades compared to Clade Ia. A dot plot showing differences across the genomes of MPXV clades in comparison to NCBI reference (NC003310, representing Clade Ia). The gaps in Clade Ib, IIa, and IIb, as compared to reference Clade Ia, are marked by circles. Clade Ib, IIa, and IIb show one, three and four major gaps, respectively. A green box demarcates the inverted terminal repeats (ITR) present on 5’ and 3’ ends of the MPXV genome.

### B22R Surface Protein Shows Amino Acid Changes Across All the MPXV Clades

MPXV appears to be altering its evolutionary trajectory by accumulating mutations as it increasingly circulates among humans. To understand these changes at the protein-level, we examined the occurrence of mutations and identified proteins with high-frequency mutations. Figure S4 (Venn diagram) illustrates the distribution of the number of SNP and SNP-silent mutations (384, 238, 815, and 874 in Clades Ia, Ib, IIa, and IIb, respectively). Figure 5A highlights the distribution of proteins showing high-frequency (>90%) non-synonymous mutations across the MPXV clades. A total of 87 high-frequency clade-specific non-synonymous mutations are distributed in three clades across the 55 MPXV proteins. MPXV_gp174 (encoding B22R glycoprotein) is commonly mutated in these three clades with 4 unique mutations in Clade IIb, followed by 2 mutations in Clade Ib and 1 mutation in Clade IIa. The highest clade-specific mutations were found in Clade IIb (43 mutations), followed by Clade IIa (26 mutations) and 18 mutations were found in Clade Ib. Notably, no unique mutations at more than 90% frequency were observed in Clade Ia (but two clade-specific mutations were observed at 50% frequency-data not shown). The presence of uniquely mutated proteins, along with certain shared mutations across clades, may indicate clade-specific differences that may contribute to variations in infection patterns and associated symptoms.

**Figure 5.**
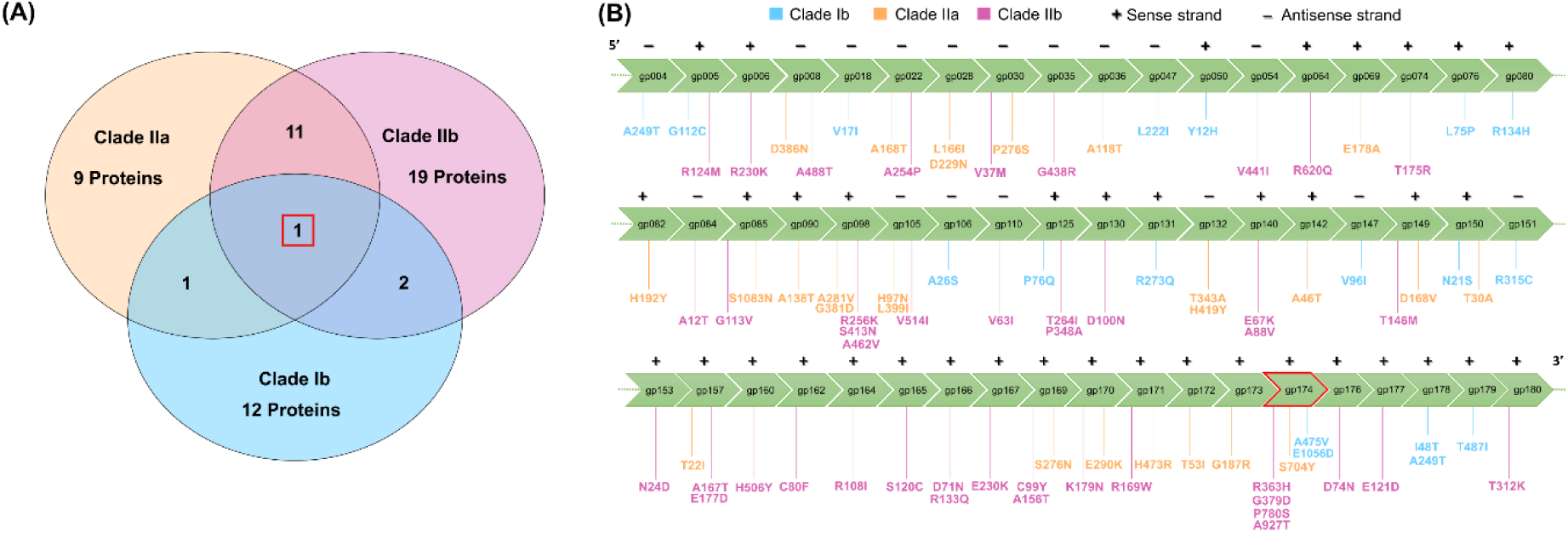
Mutational spectrum in MPXV proteins. (A) A Venn diagram illustrating uniquely mutated proteins (>90% frequency) among clades, either shared (overlapping ellipse) or specific to clades (non-overlapping region). Clade Ib (blue), Clade IIa (orange), and Clade IIb (pink) show 12, 9, and 19 uniquely mutated proteins, respectively. (B) The mutational catalogue across all the MPXV genes, with the green arrowed boxes representing the locus tags based on the MPXV NCBI reference sequence (NC_003310.1). The mutations in the MPXV locus tags are shown for Clade Ib, Clade IIa, and Clade IIb in blue, orange, and pink, respectively.

To further explore the observed differences between clades, we analysed the temporal expression and functional roles of mutated proteins using available expression data from Huang, Y. *et al.* (2024) [33]. We focused on proteins uniquely mutated in MPXV clades and their expression profiles over time. Figure S5A highlights uniquely mutated proteins in Clade Ib compared to Clade Ia. These proteins are expressed mostly at 6 hours post-infection (hpi), followed by 12 hpi and 24 hpi, with primary involvement in host immune evasion, as well as roles in replication and transcription. Figure S5B reveals 7 uniquely mutated proteins in Clade IIb and 3 in Clade IIa. In Clade IIa, all mutated proteins are expressed during the initial 6 hours, whereas in Clade IIb, 5 of the 7 proteins are expressed at 6 hpi, with 1 protein each expressed at 12 and 24 hpi. These findings highlight clade-specific differences in protein expression timing, which may be linked to their functional roles in immune evasion and viral replication.

### Ten Non-Synonymous Mutations Exhibit Damaging Effects in MPXV Virus

To assess the impact of high-frequency mutations on viral proteins, we used tools such as MutPred2 and PolyPhen2. Table 1 highlights the genotype-to-phenotype analysis of clade-specific mutations with high MutPred2 score. In Clade Ib, four proteins - EGF-like domain (MPXV_gp005), crescent membrane and immature virion formation (MPXV_gp076), hydroxysteroid dehydrogenase (MPXV_gp151), and B22R family protein (MPXV_gp174) showed MutPred2 scores of 0.799, 0.59, 0.471, and 0.419 for G112C, L75P, R315C, and E1056D mutations, respectively. These proteins are implicated in immune evasion and viral pathogenesis. In Clade IIa, four proteins displayed MutPred2 scores between 0.5 and 0.6. Notably, E178A in MPXV_gp069, and A168T in MPXV_gp022 were predicted to impact immune evasion and structural integrity. Additionally, an E290K mutation in MPXV_gp170, involved in virion assembly, showed a significant charge alteration supported by both MutPred2 and PolyPhen2. The R304C mutation in MPXV_gp008 is predicted to disrupt a critical salt bridge, leading to structural damage (data not shown). In Clade IIb, four SNP mutations - G113V (MPXV_gp085), H506Y (MPXV_gp160), C99Y (MPXV_gp169), and R169W (MPXV_gp171) - exhibited high MutPred2 scores. The C99Y mutation in MPXV_gp169 is particularly significant due to the change from cysteine, a residue involved in disulfide bond formation, to aromatic tyrosine. The analysis for other mutations is presented in Supplementary File 4. This result shows that most of the mutations are either neutral or benign, i.e., having no damaging effect on viral proteins.

**Table 1.**
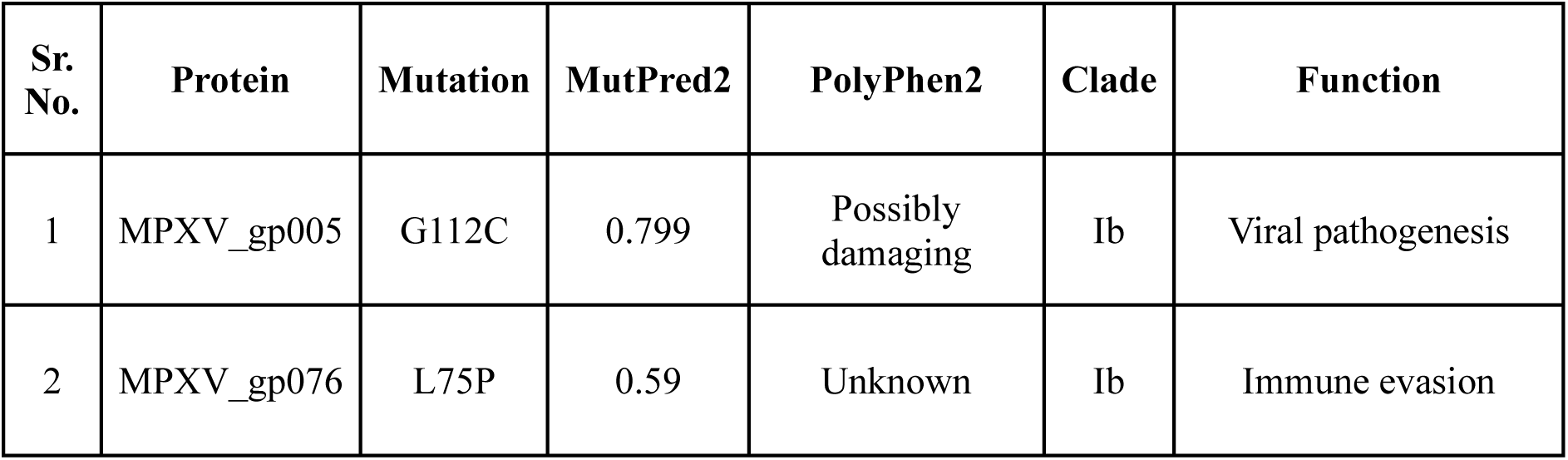
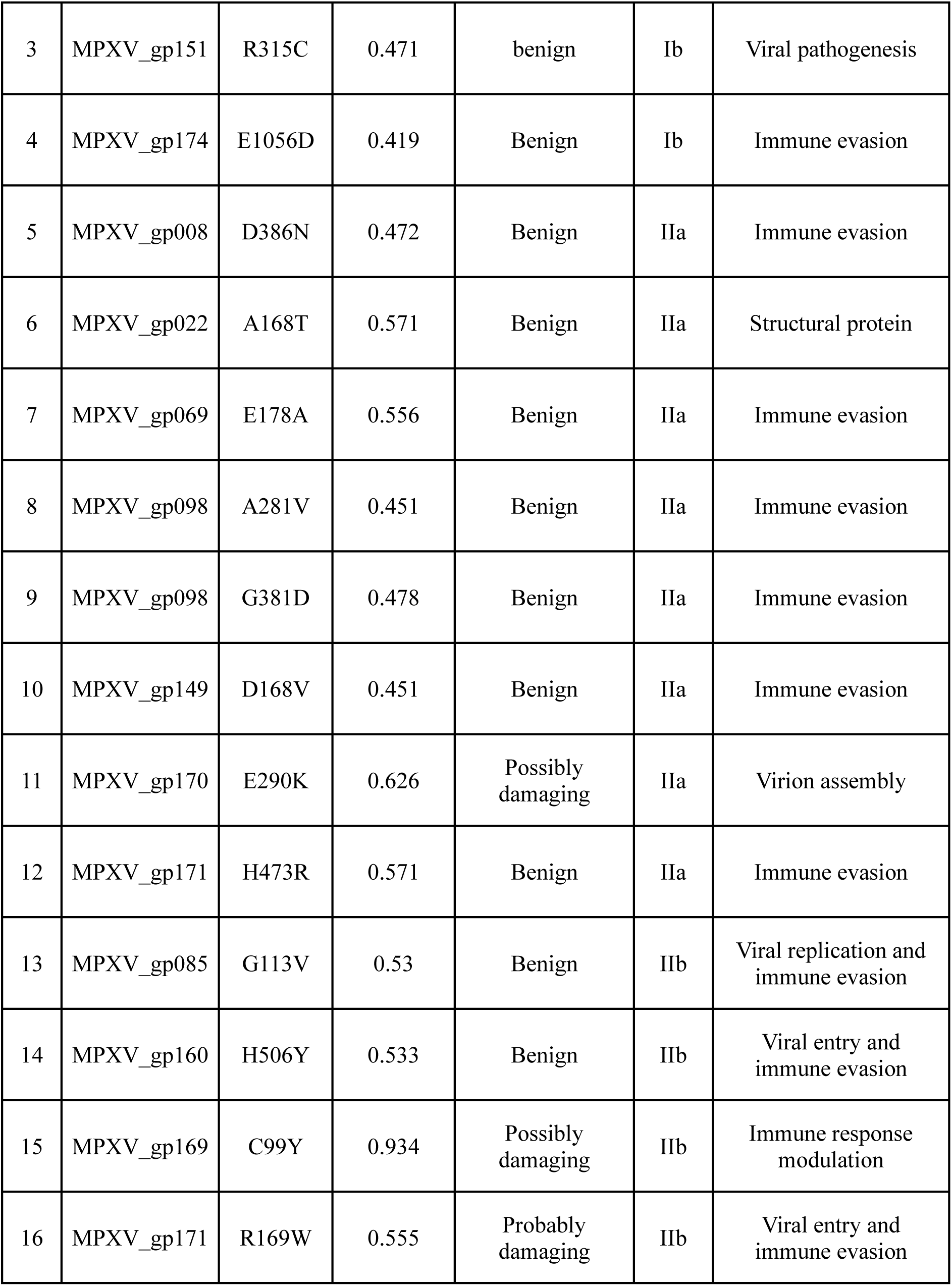
High-frequency mutations showing altered phenotype. Based on the score, the MutPred2 and PolyPhen2 analysis for 16 high-frequency mutations showed altered phenotypes. The function and the clade the high-frequency mutation is associated with is also presented.

## Discussion

The global eradication of smallpox in 1980 led to the termination of routine smallpox vaccination [34]. This vaccine is known to offer cross-protection against other orthopoxviruses, providing up to 85% protection against mpox [35, 36]. However, the cessation of smallpox vaccination left populations increasingly susceptible to other orthopoxviruses, including MPXV [36]. Previously, MPXV was primarily restricted to rodents and monkeys in Africa but has recently caused more frequent and severe outbreaks in humans [37]. In 2022, Clade IIb of MPXV caused a global outbreak with increased human-to-human transmissibility, resulting in mpox being declared a Public Health Emergency of International Concern (PHEIC) in July 2022 [4]. In 2024, another variant, Clade Ib, has spread outside Africa, raising global concern again.

There are various strategies used by viruses to evolve, such as recombination, insertions, deletions, and SNPs [38–40]. In our study, we report a 1141 bp long gap toward the 5’ end of the Clade Ib genome, indicating the deletion of CCP protein (homologous to human C4b-binding protein), which is known to counteract the host complement system. This deletion was previously considered to be specific to Clade II of MPXV [41], but we find that this deletion is present in Clade Ib sequences as well. The relevance of this deletion is unknown, as several studies have investigated the significance of CCP, and their conclusions remain inconsistent. Girgis *et al.* (2011) and Hudson *et al.* (2012) reported reduced pathogenicity upon CCP and IMP (CCP homolog in vaccinia virus)-deletion respectively [42, 43]. Interestingly, although CCP deletion caused reduced viral titres indicating less virulence, our analysis of the qPCR data from Hudson *et al*. (2012) suggests prolonged persistence of the virus. This sustained persistence may increase the chances of infection spread. On the other hand, Kotwal *et al.* (1998) showed that IMP (CCP-homolog of cowpox virus) knock-outs caused more extensive lesions in mouse footpad, concluding that higher inflammation response worsens the infection in the absence of IMP [44]. However, overlooking other routes of infection and simultaneous mutations may inevitably lead to this conundrum. Notably, Clade II, which exhibits CCP deletion, is considered to be less pathogenic whereas, Clade Ia retains an intact CCP and is found to be more pathogenic [45]. The CCP deletion in Clade Ib could potentially alter the virus’s pathogenicity compared to Clade Ia. Several other deletions and insertions were also observed in Clade IIa and Clade IIb. For instance, a gain of 4 amino acids in Ser/Thr kinase encoded by the genetic locus MPXV_gp159 for Clade IIa sequences was observed (data not shown). It is essential to acknowledge that each clade has acquired a unique set of mutations as part of its adaptive evolution.

Several orthopoxviruses infect both human and non-human hosts, while the variola virus is exclusive to humans [46–48]. The high genetic similarity among orthopoxviruses poses a continual risk to humans by zoonotic transmission and reverse zoonosis to animals (living in close proximity to humans). DNA viruses, including orthopoxviruses, are generally characterised by their low mutation rate [49]. But recent MPXV variants show increasing mutation counts and accelerated APOBEC-related nucleotide changes compared to their predecessors [32, 50]. The APOBEC family of enzymes (apolipoprotein B mRNA editing enzyme, catalytic polypeptide) is produced by all mammals, with the highest expression seen in humans [51]. APOBEC induces C>T, consequently resulting in G>A transitions on the complementary strand in viruses. Our analysis (Figure 3A) showed the elevated ratio of C>T and G>A transitions relative to all other transitions and transversions in Clade Ib as compared to Clade Ia. A similar trend of nucleotide change was observed in Clade IIb during the 2022 outbreak. These results suggest that Clade Ib may follow a similar trajectory as Clade IIb, potentially leading to a higher rate of human-to-human transmission. In contrast, Clades Ia and IIa show low C>T and G>A transitions and hence are considered to be mostly zoonotic, as supported by epidemiological data [52].

The data from genotype-to-phenotype prediction presents that most of the mutations showed no damaging effect on the viral proteins. This aligns with a previous study stating that most of the mutations led to protein stabilisation in MPXV [53]. The mutation analysis and gene expression data revealed that most mutated proteins in MPXV are involved in immune evasion, viral transcription and host cytoskeleton modification. Most of these proteins are expressed early in the virus’s life cycle, i.e., at 6- and 12-hours post-infection. MPXV_gp174 coding for the B22R family protein, which is a significant epitope and host receptor-binding protein in MPXV [54], shows high-frequency mutations like S704Y specific to Clade IIa; A927T, G379D, P780S, and R363H mutations in Clade IIb; and, A475V and E1056D mutations in Clade Ib. These mutations in this highly immunogenic protein may affect the binding affinity, virulence and transmissibility of the virus.

In conclusion, our study highlights significant genomic differences among MPXV clades and identifies high-frequency mutations that prompt further experimental investigation to understand their impact on infection mechanisms and disease progression. These mutations are crucial for developing rapid diagnostic tests, which can help control outbreaks in their early stages by enabling targeted responses. We emphasise the importance of ongoing genomic surveillance to stay ahead of the virus and enhance our overall understanding of its evolution and pathogenicity.

## Acknowledgements

The authors thank Utpal Tatu lab members for their discussions and input.

## Author contributions

Conceptualisation: AK, UT. Data Curation: AK, PJ, RB. Formal Analysis: AK, PJ, RB. Investigation: AK, PJ, RB, AG, UT. Methodology: AK, PJ, RB, UT. Supervision: UT. Project Administration: UT. Resources: UT. Validation: AK, PJ, RB, UT. Writing – Original Draft: AK, PJ, RB. Writing – Review & Editing: AK, PJ, RB, UT.

## Data Availability

The data supporting the findings in the study is available in the manuscript and as supporting data.

## Ethics approval

Not applicable

## Supplementary data

Supplementary figures and tables

Supplementary file 1. An R script to annotate SNPs for ambipolar genomes.

Supplementary file 2. An R script for generating high-frequency mutations.

Supplementary file 3. A gff3 file with coordinates for coding regions of MPXV.

Supplementary file 4. A Genotype-to-Phenotype data for high-frequency mutations found in MPXV clades

