## Supplementary material for "Genomic analyses of recently emerging clades of mpox virus reveal gene deletion and single nucleotide polymorphisms that correlate with altered virulence and transmission": Supp_File1_R_script_annotation_and_merge.docx

Authors: Original Script - Mercatelli et al. (2020)

Modified Script – Ankeet Kumar

#Bash part

ref=NC_003310.fasta

input=input.fasta

nucmer --forward -p nucmer $ref $input

show-coords -r -c -l nucmer.delta > nucmer.coords

show-snps nucmer.delta -T -l > nucmer.snps

\# Load Data (Output from nucmer- Bash)

nucmer <- read.delim("nucmer.snps", as.is = TRUE, skip = 4, header = FALSE, sep = "\t")

colnames(nucmer) <- c("refpos", "refvar", "qvar", "qpos", "", "", "", "", "rlength", "qlength", "", "", "rname", "qname")

rownames(nucmer) <- paste0("var", 1:nrow(nucmer))

\# Fix IUPAC codes

table(nucmer$qvar)

nucmer<-nucmer[!nucmer$qvar%in%c("B","D","H","K","M","N","R","S","V","W","Y"),]

nrow(nucmer)

\### Aminoacid variant list ----

\# Load reference sequence

library(seqinr)

library(Biostrings)

refseq<-read.fasta("NC_003310.fasta",forceDNAtolower=FALSE)[[1]]

\# Load GFF3

gff3<-read.delim("NC_003310.gff3",as.is=TRUE,skip=5,header=FALSE)

annot<-setNames(gff3[,10],gff3[,9])

header<-c("sample","refpos","refvar","qvar","qpos","qlength","protein","variant","varclass","annotation")

results<-matrix(NA,ncol=length(header),nrow=0)

colnames(results)<-header

samples<-unique(nucmer$qname)

pb<-txtProgressBar(0,length(samples),style=3)

for (pbi in seq_along(samples)) {

sample <- samples[pbi]

allvars <- nucmer[nucmer$qname == sample, ]

for (i in seq_len(nrow(allvars))) {

nucline <- allvars[i, ]

refpos <- as.numeric(nucline["refpos"])

refvar <- nucline["refvar"]

qvar <- nucline["qvar"]

qpos <- as.numeric(nucline["qpos"])

qlength <- nucline["qlength"]

\# Match over GFF3 annotation

a <- refpos - gff3[, 4]

b <- refpos - gff3[, 5]

signs <- sign(a) * sign(b)

w <- which(signs == -1)

\# Initialize default values

protein <- "extragenic"

output <- c(refpos, "extragenic", "UTR")

if (length(w) == 0) {

\# Extragenic regions

if (refpos < gff3[1, 4]) {

protein <- "5'UTR"

} else if (refpos > gff3[nrow(gff3), 5]) {

protein <- "3'UTR"

} else {

protein <- "intergenic"

}

results <- rbind(results, data.frame(

sample = sample,

refpos = refpos,

refvar = refvar,

qvar = qvar,

qpos = qpos,

qlength = qlength,

protein = protein,

variant = output[1],

varclass = output[2],

annotation = output[3],

stringsAsFactors = FALSE

))

}

else {

\# Within gene

for (j in seq_along(w)) {

start <- gff3[w[j], 4]

end <- gff3[w[j], 5]

protein <- gff3[w[j],9]

annotation<- gff3[w[j],10]

current_strand <- gff3[w[j], 7]

if (qvar == ".") {

\# Handle deletions

if ((nchar(refvar) %% 3) != 0) {

if (current_strand == "-") {

mutpos <- ceiling((end - refpos) / 3)+1

} else {

mutpos <- ceiling(((refpos + 1) - start) / 3)

}

output <- c(paste0(refpepseq[mutpos], mutpos), "deletion_frameshift")

} else {

varseq <- refseq[-(refpos:(refpos + nchar(refvar) - 1))]

varseq <- varseq[start:(end - nchar(refvar))]

vardnaseq <- Biostrings::DNAString(paste0(varseq, collapse = ""))

varpepseq <- Biostrings::translate(vardnaseq)

varpepseq <- strsplit(as.character(varpepseq), "")[[1]]

for (k in seq_along(refpepseq)) {

if (refpepseq[k] != varpepseq[k]) {

output <- c(paste0(refpepseq[k], k), ifelse(varpepseq[k] == "*", "deletion_stop", "deletion"))

break

}

}

}

} else if (refvar == ".") {

\# Handle insertions

if ((nchar(qvar) %% 3) != 0) {

if (current_strand == "-") {

mutpos <- ceiling((end - refpos) / 3)+1

} else {

mutpos <- ceiling(((refpos + 1) - start) / 3)

}

output <- c(paste0(refpepseq[mutpos], mutpos), "insertion_frameshift")

} else {

varseq <- c(refseq[1:refpos], strsplit(qvar, "")[[1]], refseq[(refpos + 1):length(refseq)])

vardnaseq <- Biostrings::DNAString(paste0(varseq[start:(end + nchar(qvar))], collapse = ""))

varpepseq <- Biostrings::translate(vardnaseq)

varpepseq <- strsplit(as.character(varpepseq), "")[[1]]

for (k in seq_along(refpepseq)) {

if (refpepseq[k] != varpepseq[k]) {

nr_aa_inserted <- nchar(qvar) / 3

multivarj <- varpepseq[k:(k + nr_aa_inserted - 1)]

output <- c(paste0(paste0(multivarj, collapse = ""), k), ifelse(any(multivarj == "*"), "insertion_stop", "insertion"))

break

}

}

}

} else if (nchar(qvar) == 1) {

\# Handle SNPs

if (current_strand == "-") {

codonpos <- ceiling((end - refpos) / 3)+1

codonstart <- (end/ 3) -2

codonend <- codonstart - 2

varseq <- refseq

varseq[refpos] <- qvar

vardnaseq <- Biostrings::DNAString(paste0(rev(comp(varseq[start:end])),collapse = ""))

varpepseq <- Biostrings::translate(vardnaseq)

varpepseq <- strsplit(as.character(varpepseq), "")[[1]]

refdnaseq <- Biostrings::DNAString(paste0(rev(comp(refseq[start:end])),collapse = ""))

refpepseq <- Biostrings::translate(refdnaseq)

refpepseq <- strsplit(as.character(refpepseq), "")[[1]]

refaa <- refpepseq[codonpos]

varaa <- varpepseq[codonpos]

output <- c(paste0(refaa, codonpos, varaa), ifelse(refaa == varaa, "SNP_silent", ifelse(varaa == "*", "SNP_stop", "SNP")))

} else {

codonpos <- ceiling(((refpos +1) - start) / 3)

codonstart <- (codonpos - 1) * 3 + start

codonend <- codonstart + 2

varseq <- refseq

varseq[refpos] <- qvar

vardnaseq <- Biostrings::DNAString(paste0(varseq[start:end], collapse = ""))

varpepseq <- Biostrings::translate(vardnaseq)

varpepseq <- strsplit(as.character(varpepseq), "")[[1]]

refdnaseq <- Biostrings::DNAString(paste0(refseq[start:end], collapse = ""))

refpepseq <- Biostrings::translate(refdnaseq)

refpepseq <- strsplit(as.character(refpepseq), "")[[1]]

refaa <- refpepseq[codonpos]

varaa <- varpepseq[codonpos]

output <- c(paste0(refaa, codonpos, varaa), ifelse(refaa == varaa, "SNP_silent", ifelse(varaa == "*", "SNP_stop", "SNP")))

}

}

\# Append results to the dataframe

results <- rbind(results, data.frame(

sample = sample,

refpos = refpos,

refvar = refvar,

qvar = qvar,

qpos = qpos,

qlength = qlength,

protein = protein,

variant = output[1],

varclass = output[2],

annotation =annotation ,

stringsAsFactors = FALSE

))

}

}

}

setTxtProgressBar(pb, pbi)

}

\# Write results to CSV

write.csv(results, "results.csv", row.names = FALSE)

\# Load the library required for merging script

library(dplyr)

\# merge the results and metadata to contain inforamtion of clades with the sequences

results1<- results%>%

mutate(qpos = as.character(qpos),

qlength = as.character(qlength),

refpos = as.character(refpos))

\# produce a dataframe of merged results

merged_results<- results1%>%

group_by(sample, variant, protein) %>%

#filter(n() <= 3) %>%

reframe(

refpos = paste(refpos, collapse = "_"),

refvar = paste(refvar, collapse = ""),

qvar = paste(qvar, collapse = ""),

qpos = paste(qpos, collapse = "_"),

qlength = first(qlength),

varclass = first(varclass),

annot = first(annotation))

write.csv(merged_results,"MERGED_RESULTS.csv", row.names = F)
