## Supplementary material for "Genomic analyses of recently emerging clades of mpox virus reveal gene deletion and single nucleotide polymorphisms that correlate with altered virulence and transmission": Supp_File2_R_Script_for_High_frequency_mutations.docx

library(dplyr)

library(tidyr)

\# Load data

merged_results<- read.csv("merged_results.csv")

variant<- merged_results

\# Remove extragenic mutations

variant <- subset(variant, varclass!= "extragenic")

\# Merge two coloumns so that if same variant is present in two columns it doesn't hamper your results

variant$protvar <- paste(variant$protein, variant$variant,sep = "_")

variant%>% group_by(clade)%>%summarize(Total= n_distinct(sample))

\# Setup a cutoff for minimum samples used

cutoff<-variant%>% group_by(clade)%>%summarize(Total= n_distinct(sample)) %>% filter(Total>=4)

\# Check clades that pass the cutoff

cutoffclade<- cutoff$clade

\# Remove clades that do not pass threshold

variant<-variant[variant$clade %in% cutoffclade,]

\# Inititate a list

mutation_list <- list()

\# Get unique major clades

clades <- unique(variant$clade)

\# Iterate through each major clade

for (clade in clades) {

\# Subset the data for the current major clade

clade_data <- variant[variant$clade == clade, ]

\# Get the mutations for the current major clade

mutations <- clade_data$protvar

\# Add the mutations to the list with the major clade name as the list name

mutation_list[[clade]] <- mutations

}

\# Print the mutation list

mutation_list

#find_common variants

common_variants <- Reduce(intersect, mutation_list)

common_variants

clade_pairs <- combn(names(mutation_list), 2, simplify = FALSE)

\# Shared mut

if(length(clade_pairs) >=2) {

clade_pairs <- combn(names(mutation_list), 2, simplify = FALSE)

\# Iterate through each clade pair

for (pair in clade_pairs) {

\# Get the clade names

clade1 <- pair[1]

clade2 <- pair[2]

\# Find the shared elements between the two clades

shared_elements <- intersect(mutation_list[[clade1]], mutation_list[[clade2]])

\# Print the shared elements with clade pair name

cat(paste(clade1, clade2, sep = "_"), "\n")

print(shared_elements)

cat("\n")

}

}

\# clade specific mutations

\# Set names of clade sets

names(mutation_list) <- unique(variant$clade)

\# Identify the common variants for all clades

common_variants <- Reduce(intersect, mutation_list)

\# Create a list of the unique variant sets for each clade

unique_sets <- lapply(mutation_list, function(x) setdiff(x, common_variants))

common_pairs <- list()

for (i in 1:(length(mutation_list)-1)) {

for (j in (i+1):length(mutation_list)) {

common_variants_pair <- intersect(mutation_list[[i]], mutation_list[[j]])

if (length(common_variants_pair) > 0) {

common_pairs[[paste(names(mutation_list)[i], names(mutation_list)[j], sep="_")]] <- common_variants_pair

}

}

}

non_overlapping_sets <- list()

for (i in 1:length(mutation_list)) {

overlapping_variants <- c()

for (j in 1:length(common_pairs)) {

if (names(common_pairs)[j] %in% names(mutation_list)[i]) {

overlapping_variants <- union(overlapping_variants, common_pairs[[j]])

}

}

non_overlapping_sets[[names(mutation_list)[i]]] <- setdiff(unique_sets[[i]], overlapping_variants)

}

unique_values <- lapply(non_overlapping_sets, function(x) setdiff(x, unlist(common_pairs)))

charuniq<-unique(unlist(unique_values))

clade_protvar_freq <- variant %>%

group_by(clade, protvar) %>%

filter(protvar %in% common_variants) %>%

summarise(freq = n()) %>%

mutate(max_freq = max(freq)) %>%

filter(freq >= 0.9 * max_freq) %>%

ungroup()

\# Create a pivot table to get mutation frequencies for each clade

mutation_freq_table <- pivot_wider(clade_protvar_freq, names_from = clade, values_from = freq, names_prefix = "freq_")

\# Rename columns to remove "freq_" prefix

colnames(mutation_freq_table) <- gsub("^freq_", "", colnames(mutation_freq_table))

\# Fill missing values with 0

mutation_freq_table[is.na(mutation_freq_table)] <- 0

#write.csv(mutation_freq_table, "clade_mutation_freq_table_unique.csv", row.names = F)

#{r clade_specif_mut >90%}

count_clade <- variant %>%

group_by(clade) %>%

summarise(Count = n_distinct(sample))

clade_protvar_freq <- variant %>%

filter(varclass=="SNP")%>%

group_by(clade, protvar) %>%

filter(protvar %in% charuniq) %>%

summarise(freq = n()) %>%

left_join(count_clade, by = "clade") %>%

mutate(max_freq = max(Count)) %>%

filter(freq >= 0.9 * max_freq) %>%

ungroup()

clade_protvar_freq1 <- clade_protvar_freq %>%

mutate(protein = sub("(_[^_]*_).*", "\\1", protvar))

unique_proteins <- clade_protvar_freq1 %>%

group_by(clade) %>%

#reframe(unique_proteins = paste(unique(protein), sep = ","))

reframe(unique_proteins = list(unique(protein)))

list_columns <- sapply(unique_proteins, is.list)

\# Convert list columns to character

unique_proteins[list_columns] <- lapply(unique_proteins[list_columns], function(x) sapply(x, toString))

\# Write to a file

write.table(unique_proteins, "unique_clspecficic_proteins_high_freq.tsv", sep = "\t", row.names = FALSE, quote = FALSE)

clade_protvar_freq1 <- clade_protvar_freq %>%

mutate(protein = sub("^[^_]*_[^_]*_", "", protvar))

variants_clades <- clade_protvar_freq1 %>%

group_by(clade) %>%

#reframe(unique_proteins = paste(unique(protein), sep = ","))

reframe(variants_clades = list(unique(protein)))

\#

\# unique_variations <- clade_protvar_freq %>%

\#	distinct(protvar) %>%

\#	mutate(variation = sub("^[^_]*_[^_]*_", "", protvar))

clade_protvar_freq1 <- clade_protvar_freq %>%

left_join(unique_variations, by = "protvar")

list_columns <- sapply(variants_clades, is.list)

\# Convert list columns to character

variants_clades[list_columns] <- lapply(variants_clades[list_columns], function(x) sapply(x, toString))

\# Write to a file

write.table(variants_clades, "variants_clades_high_freq.tsv", sep = "\t", row.names = FALSE, quote = FALSE)

mutation_count <- merged_results %>%

group_by(protein,clade) %>%

summarise(total = n()) %>%

left_join(count_clade, by = "clade") %>%

left_join(gff3)%>%

mutate(normalized_count= total/length)

write.csv(mutation_count, "clade_mutation_count.csv")

\# Create a pivot table to get mutation frequencies for each clade

mutation_freq_table <- pivot_wider(clade_protvar_freq, names_from = clade, values_from = freq, names_prefix = "freq_")

\# Rename columns to remove "freq_" prefix

colnames(mutation_freq_table) <- gsub("^freq_", "", colnames(mutation_freq_table))

\# Fill missing values with 0

mutation_freq_table[is.na(mutation_freq_table)] <- 0

write.csv(mutation_freq_table, "clade_specific_90percent_SNP_mutations.csv", row.names = F)

#{r univeral mut}

\# Find the common variants in the first clade

common_mutations <- Reduce(intersect,mutation_list)

\# common_variants_all will now contain the variants that are present in all clades and not shared with other clades

max_freq <- variant %>%

group_by(clade) %>%

summarise(max_freq = n_distinct(sample))

universal_protvar_freq <- variant %>%

filter(protvar %in% common_mutations)

Universalmut<- universal_protvar_freq%>%

group_by(clade, protvar, protein, varclass) %>%

summarise(freq = n()) %>%

left_join(max_freq, by = "clade") %>%

filter(freq >= 0.9 * max_freq)

#write.csv(universal_protvar_freq, "clade_Universal_mutations_all.csv", row.names = F)

mutation_listuni <- list()

\# Get unique major clades

clades <- unique(Universalmut$clade)

\# Iterate through each major clade

for (clade in clades) {

\# Subset the data for the current major clade

proteindat <- Universalmut[Universalmut$clade == clade, ]

\# Get the mutations for the current major clade

mutations <- proteindat$protvar

\# Add the mutations to the list with the major clade name as the list name

mutation_listuni[[clade]] <- mutations

}

\# Print the mutation list

mutation_listuni

common_mutationsuni <- Reduce(intersect,mutation_listuni)

universal_protvar_freq1 <- variant %>%

filter(protvar %in% common_mutationsuni)

Universalmut1<- universal_protvar_freq1%>%

group_by(clade, protvar, protein, varclass) %>%

summarise(freq = n()) %>%

left_join(max_freq, by = "clade") %>%

filter(freq >= 0.9 * max_freq)

write.csv(Universalmut1, "Universal_mutations.csv", row.names = F)
