## Supplementary material for "Genomic analyses of recently emerging clades of mpox virus reveal gene deletion and single nucleotide polymorphisms that correlate with altered virulence and transmission": Supplementary_Figures_And_Table.pdf

Affiliation:

**Figure S1. Global Overview of Mpox Disease Data.** A. Reported cases of Mpox across eight regions and worldwide from 2022 to 2024, highlighting trends in infection rates over three years. B. Detailed timeline on Mpox outbreaks since the discovery of the virus

**(A)**

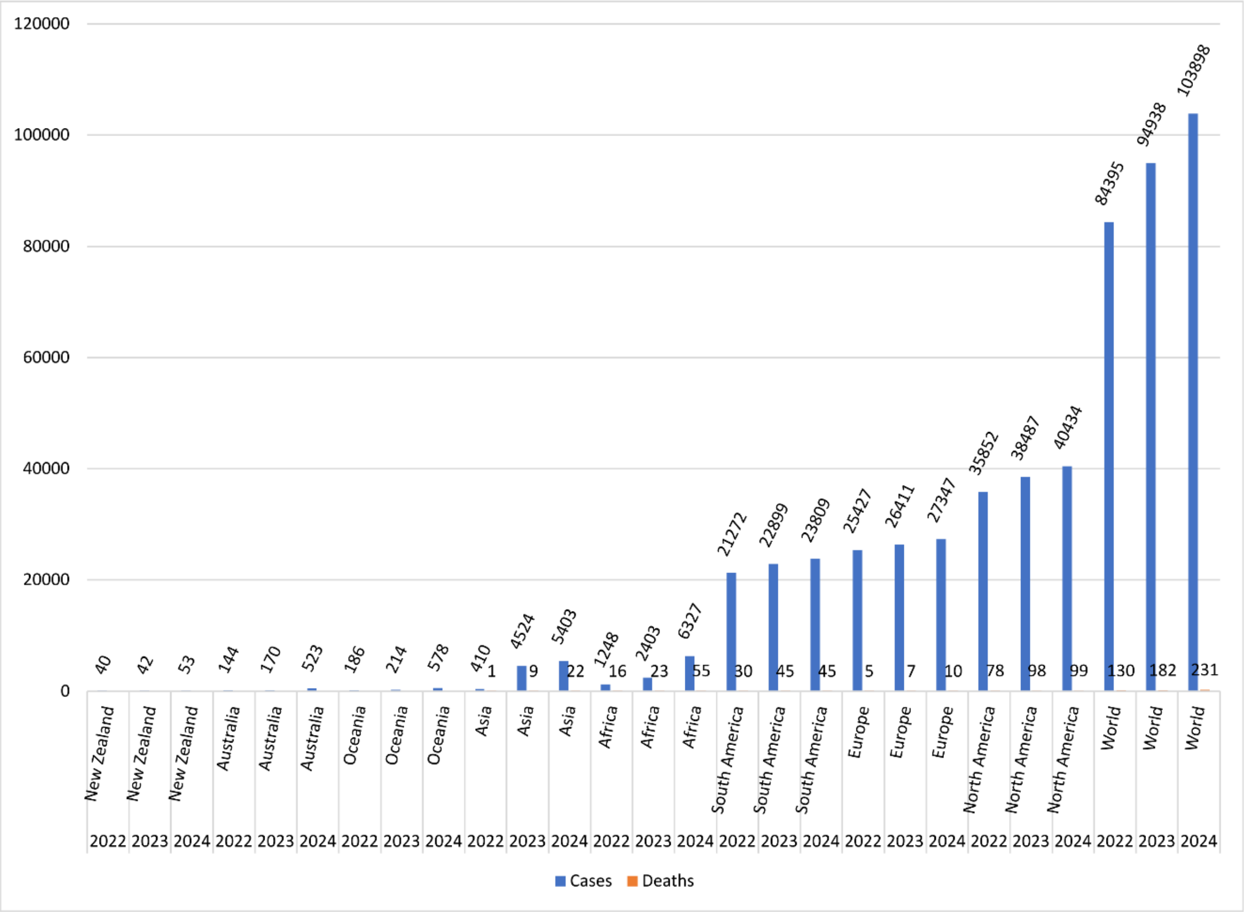


**(B)**

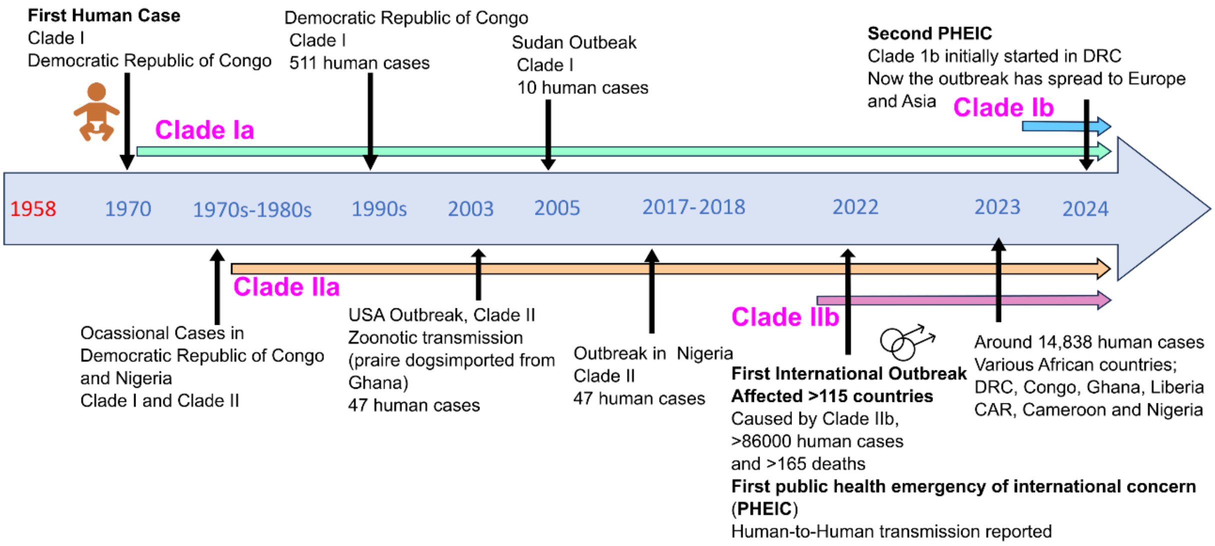

**Figure S2. Nucleotide Transition and Transversion Analysis Across Four MPXV Clades.** This figure presents the distribution of nucleotide transitions and transversions across the genome for four distinct MPXV clades. Data is based on an analysis of 10 representative sequences from each clade, providing insight into mutation patterns throughout the genome.

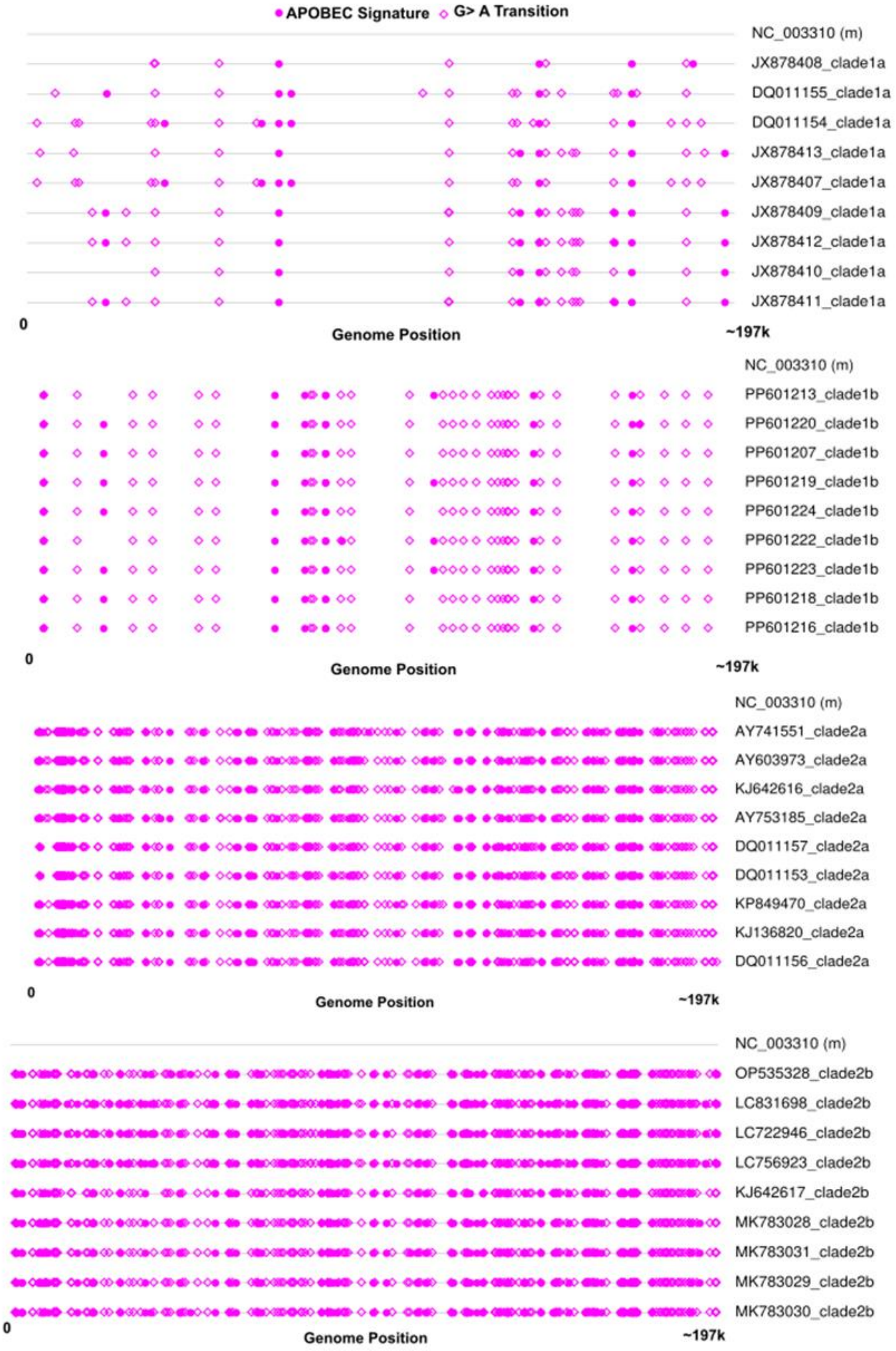

**Figure S3. CCP Gene Deletion in Clade Ib MPXV Sequences**. This figure illustrates an 1141 bp deletion in the CCP gene sequences compared to the reference sequence (NC_003310). The CCP gene is highlighted in blue.

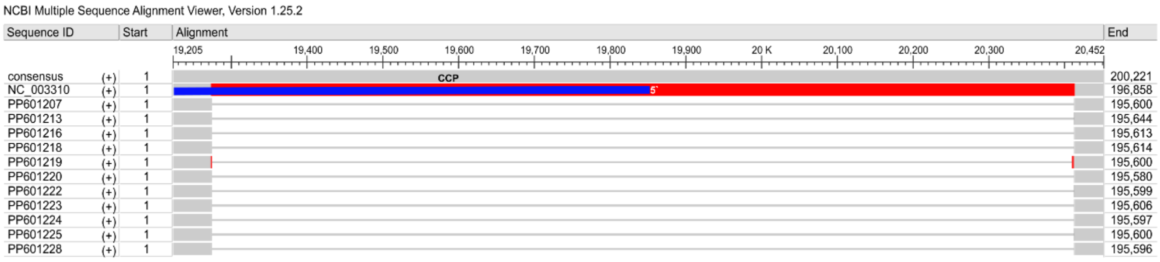


**Figure S4. Venn Diagram of Synonymous and Non-Synonymous Mutations Across Four MPXV Clades.** This figure depicts the overlap and distribution of synonymous and non-synonymous amino acid changes among four MPXV clades. The shared mutated are present in overlapping regions, and unique mutations are in non-overlapping regions.

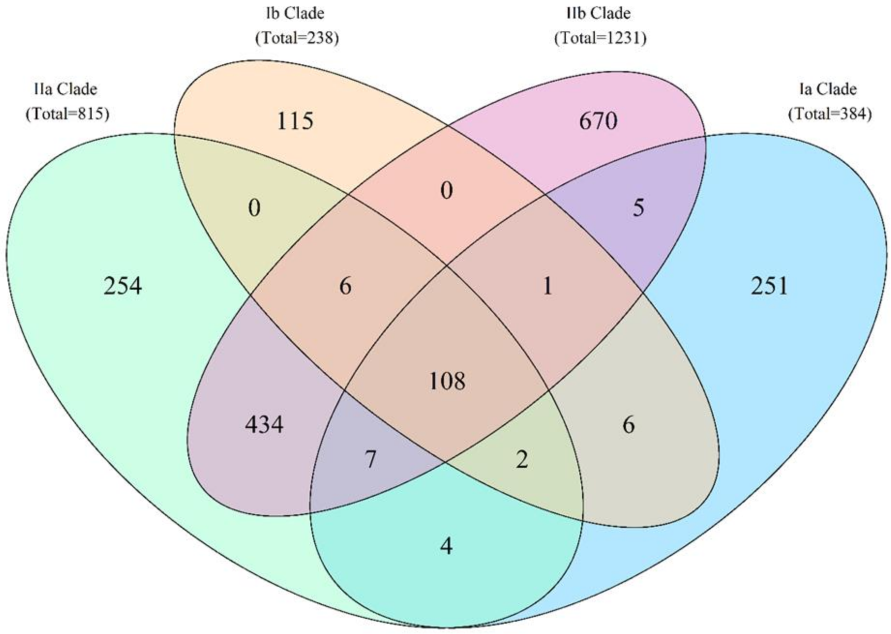

**Figure S5. Mutated proteins and Temporal Expression in MPXV Clades.** This figure illustrates the proteins affected by mutations in MPXV and their expression at different time points post-infection. Clade Ib shows more mutated proteins, with those expressed at 6 hours post-infection (hpi) being the most affected. In Clade IIb, fewer proteins are mutated; similarly, those expressed at 6 hpi are the most impacted. Notably, EGF-like domain protein and IMV membrane protein are commonly mutated in both Clade Ib and Clade IIb.

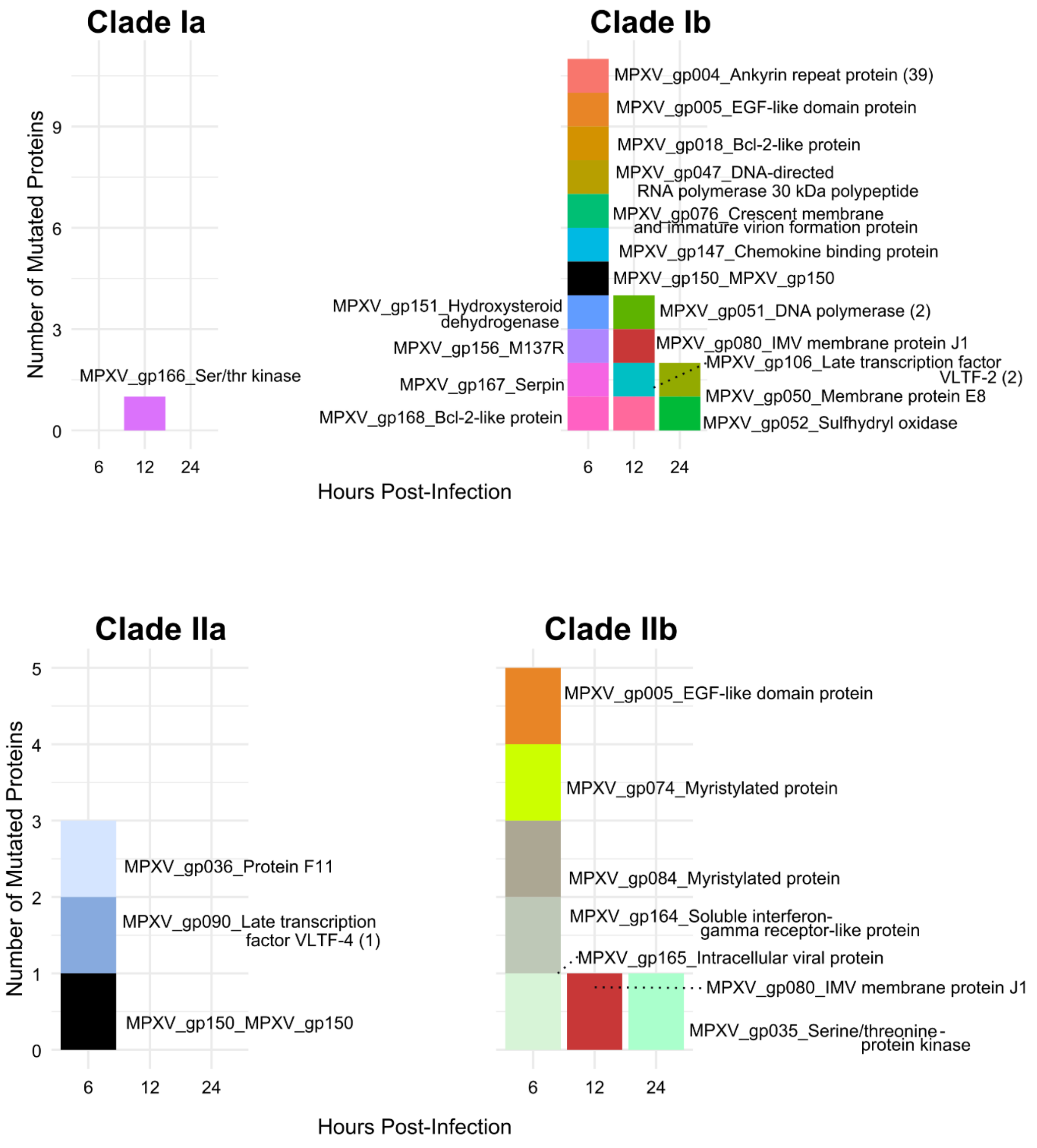

**Table S1. Nucleotide Similarity and Identity Comparison Between MPXV and Other Orthopoxviruses, Including Inter-clade MPXV Comparisons.** This table compares nucleotide similarity and identity between MPXV and other orthopoxviruses and among different MPXV clades.

| **Sequence 1** | **Sequence 2** | **Identity** | **Similarity** | **Gaps** |
|---|---|---|---|---|
| Clade Ia (NC_003310) | Clade Ib (PP601207) | 98.90% | 98.90% | 0.70% |
| Clade Ia (NC_003310) | Clade IIa (AY603973) | 95.40% | 95.40% | 4.10% |
| Clade Ia (NC_003310) | Clade IIb (OP535328) | 96.50% | 96.50% | 3.00% |
| Clade Ib(PP601207) | Clade IIa (AY603973) | 95.80% | 95.80% | 3.50% |
| Clade Ib (PP601207) | Clade IIb (OP535328) | 97.00% | 97.00% | 2.40% |
| Clade IIa (AY603973) | Clade IIb (OP535328) | 98.30% | 98.30% | 1.40% |
| MPXV (NC_003310) | ABMPV (MH816996) | 85.70% | 85.70% | 9.41% |
| MPXV (NC_003310) | AKHV (MH607141) | 80.47% | 80.47% | 12.78% |
| MPXV (NC_003310) | CMPV (AF438165) | 84.47% | 84.47% | 11.28% |
| MPXV (NC_003310) | CPV (AF482758) | 82.11% | 82.11% | 11.28% |
| MPXV (NC_003310) | ECTV (AF012825) | 82.70% | 82.70% | 12.00% |
| MPXV (NC_003310) | RAPV (KP143769) | 72.80% | 72.80% | 11.60% |
| MPXV (NC_003310) | SKPV (KU749310) | 71.30% | 71.30% | 14.00% |
| MPXV (NC_003310) | TATPV (DQ437594) | 87.10% | 87.10% | 8.70% |
| MPXV (NC_003310) | VACV (AY243312) | 86.20% | 86.20% | 8.50% |
| MPXV (NC_003310) | VARV (X69198) | 84.50% | 84.50% | 11.20% |

| | | | | |
|---|---|---|---|---|
| MPXV (NC_003310) | VPXV (KU749311) | 71.70% | 71.70% | 12.80% |
